# A cohesin-centric gene regulatory network resource with regulatory site annotations

**DOI:** 10.64898/2026.01.06.697865

**Authors:** Jinyu Ding, Haoping Chen, Yanqiang Fu, Wenze Gao, Zhixiong Fang, Jiankang Wang

## Abstract

Cohesin is a central regulator of transcription and chromatin organization, binding to the majority of cis-regulatory elements (CREs) and mediating enhancer-promoter communication through three-dimensional genome architecture. Although gene regulatory networks (GRNs) provide an interpretable framework for modeling transcriptional regulation, most existing GRN analyses focus on transcription factor (TF)-gene relationships and largely ignore regulatory sites. Consequently, cohesin-associated regulatory contexts are rarely incorporated into GRN reconstruction. Here, we present a cohesin-centric gene regulatory network database that explicitly integrates TF binding, regulatory sites, and gene targets into unified TF-site-gene regulatory paths. Building upon our previously developed multiomics resource CohesinDB, we mapped TF binding to cohesin-associated regulatory sites and linked these sites to their downstream target genes. The resulting Cohesin-GRN module in CohesinDB (http://cohesindb.wangjklab.com/) (http://120.24.147.32/)comprises 61,222,502 TF-cohesin site links and 2,228,634 cohesin site-gene links, collectively forming over 270 million TF-site-gene regulatory paths. By enabling a CRE-informed and site-aware representation of gene regulation, Cohesin-GRN bridges conventional TF-gene GRNs with regulatory site-centric mechanisms. Given the pervasive roles of cohesin in enhancer activity, transcriptional control, and human disease, Cohesin-GRN provides a valuable resource for exploring transcriptional dysregulation and gene regulatory networks.

## Introduction

Cohesin is a ring-shaped chromatin-binding protein complex composed of multiple subunits, including Rad21, SMC1, SMC3, and SA1/2 [1]. Unlike early studies that primarily focused on its canonical role in holding sister chromatids together during mitosis, modern molecular biology has established cohesin as a central regulator of transcriptional control and chromatin folding [2]. Cohesin is involved in nearly all enhancer activities [3] and directly influences gene expression [4]. In mammalian cells, cohesin extrudes chromatin through a “loop extrusion” mechanism [1] and, upon encountering the insulator protein CTCF, contributes to the formation of higher-order chromatin structures such as topologically associated domains. Cohesin can also function independently of CTCF by mediating cis-regulatory elements (CREs) enriched with transcription factors (TFs) and epigenetic modifications, thereby promoting chromatin loop formation between enhancers and promoters [1]. Consequently, transcription critically depends on the precise regulation of chromatin three-dimensional architecture and cis-regulatory elements by cohesin [2, 5, 6]. Dysregulation of cohesin disrupts chromatin folding and transcriptional homeostasis, leading to developmental abnormalities, genetic disorders, cancer, and a wide range of human diseases [4, 6, 7]. For example, mutations in cohesin-related genes such as NIPBL and SMC1 cause Cornelia de Lange syndrome (CdLS), a congenital developmental disorder [2, 7, 8]. Moreover, cohesin is among the most frequently mutated gene complexes in cancer, acting as a driver in multiple tumor types [4, 9]. In recent years, cohesin-related research has emerged as a major entry point for understanding transcriptional regulation, three-dimensional genome organization, and the molecular basis of human disease [1-6, 8-16].

On the other hand, gene expression is tightly regulated by complex networks of TFs and cofactors [17]. Deciphering these networks is a central goal of modern computational biology [18]. The interplay between chromatin, TFs and genes generates complex regulatory circuits that can be represented as gene regulatory networks (GRNs) [19]. GRNs are interpretable computational representations of gene expression regulation, modeled as networks in which nodes correspond to genes and edges represent regulatory interactions, primarily between TFs and their target genes [20]. GRNs capture how combinatorial TF activity governs transcriptional programs underlying cellular function. Elucidating the topology and dynamics of GRNs is fundamental to understanding the establishment and maintenance of cellular identity, with important implications for cell fate control and disease mechanisms. However, transcriptional regulation is not directly mediated by TF–gene interactions. Instead, regulatory control is exerted through CREs and chromatin architecture, indicating that accurate GRN reconstruction requires integrating regulatory contexts [21, 22]. CREs, including promoters, enhancers, and other distal regulatory sites, play a central role in gene regulation by mediating the physical and functional interactions between TFs and their target genes [23]. Acting as the primary substrates of transcriptional regulation, CREs integrate multiple epigenetic cues to fine-tune gene expression programs [24].

Importantly, accumulating evidence indicates that cohesin binds to the majority of active CREs and is critically involved in organizing enhancer–promoter communication through chromatin looping and higher-order genome architecture [18-24]. Despite the recognized importance of CREs and cohesin in transcriptional regulation [11-15, 25-27], most existing GRN models are constructed without explicitly incorporating cohesin-associated regulatory context. Consequently, current GRN analyses often overlook the cohesin-mediated chromatin interactions, limiting their ability to represent more regulatory information. Previously, we developed CohesinDB [14, 15], a comprehensive cohesin-centered database that integrates epigenomes, 3D genomes and transcriptomes in human cells. Building upon this foundation, we systematically mapped transcription factor (TF) binding to cohesin-associated regulatory sites and linked these cohesin sites to their downstream target genes. Based on this TF–site–gene framework, we constructed Cohesin-GRN, a cohesin-annotated gene regulatory network that explicitly incorporates regulatory site information. This new module of CohesinDB (Cohesin-GRN) comprises 61,222,502 TF-cohesin site links and 2,228,634 cohesin site-gene links, collectively forming over 270 million TF– site–gene regulatory paths. Given that cohesin participates in nearly all enhancer activities and directly regulates gene expression [1,6], Cohesin-GRN provides a broadly applicable resource for studying transcriptional regulation and gene regulatory network architecture.

## Methods

### Database design

In this study, we constructed and implemented Cohesin-GRN, an online query and visualization platform dedicated to cohesin-associated gene regulatory networks. Cohesin-GRN is developed as an integrated functional module within the CohesinDB framework. The system adopts a standard web-based service architecture, with MySQL (version 8.0.44) serving as the core relational database management system and Django (version 3.0.14) used as the primary framework for both backend and frontend development. As illustrated in **Figure 1**, the overall architecture supports a complete workflow encompassing data storage, query execution, result integration, and interactive visualization. The backend is implemented in Python (version 3.7.16) using the Django framework, while MySQL is employed for efficient storage and management of large-scale regulatory network data. The frontend interface is developed using HTML, CSS, and JavaScript, with jQuery (version 3.6.0) facilitating dynamic content interaction and Bootstrap (version 5.3.0) enabling responsive web design across different devices. Cohesin-GRN is deployed on an Ubuntu-based (version 24.04) server environment using Nginx (version 1.24.0) as the web server. The database is freely accessible at http://cohesindb.wangjklab.com without login requirements and is compatible with all major modern web browsers (e.g., Google Chrome), provided that JavaScript is enabled.

**Figure 1.**
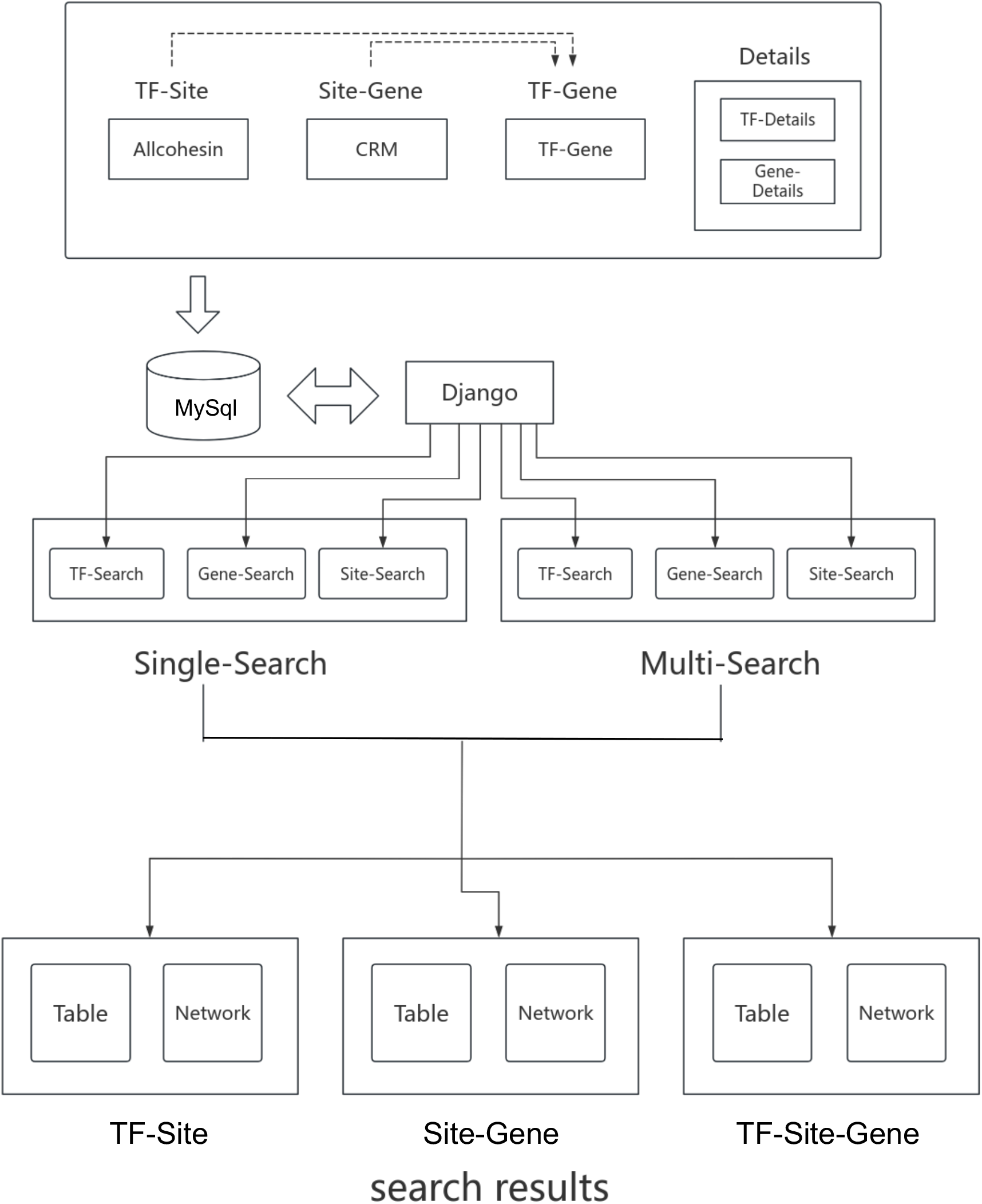
Construction and overall architecture of the Cohesin-GRN database. Schematic overview of data sources, network construction, database organization, and web-based query and visualization workflow implemented in Cohesin-GRN.

### Data for constructing network

Cohesin-GRN integrates multiple complementary resources. Specifically, the ‘all cohesin’ database consists of 751,590 cohesin binding sites which represent cis-regulatory elements as described in our previous studies [14, 15]. It is primarily used to store association information between TFs and regulatory sites, constituting TF–site relationships. Meanwhile, the ‘cis-regulatory module’ database records correspondences between regulatory sites and their target genes and is used to construct site–gene relationships. It was established in our previous work and contains 2,229,500 double-evidenced cis-regulatory modules. Based on these two data sources, we inferred TF–gene regulatory relationships through shared regulatory sites, thereby establishing a complete three-layer TF–site–gene regulatory network. Furthermore, Cohesin-GRN incorporates additional annotation resources, including cohesin site details derived from [15], TF details derived from [28]), and gene details derived from [29]), to provide functional annotations for regulatory sites, TFs, and target genes.

### Web Interface

At the application level, Cohesin-GRN provides flexible search functionality, supporting both single-search and multi-search modes. In the single-search mode, each query allows the input of one TF, one gene, one regulatory site, or any combination thereof. The multi-search mode enables users to input multiple TFs, genes, and sites simultaneously within a single query, thereby facilitating batch and combinatorial searches. Both search modes support arbitrary combinations of TF, gene, and site queries to accommodate diverse analytical scenarios. At the result presentation level, summary statistics of the query results are displayed as bar charts using the ECharts visualization library. Cohesin-GRN always return three categories of regulatory relationships: TF–site, site–gene, and TF–site–gene. Each category is visualized using both tabular and graphical formats, including downloadable tables implemented with DataTables (version 1.11.5) and interactive force-directed network graphs generated using ECharts, allowing intuitive exploration of the regulatory data. When browsing query results in the displayed tables, users can further interact with individual regulatory sites, TFs, or genes to access their corresponding functional descriptions and annotations.

## Results

### Overall statistics of the cohesin-centric gene regulatory network

The overall scale and composition of the cohesin-centered gene regulatory network implemented in Cohesin-GRN are summarized in **Figure 2A**. All counts are displayed on a logarithmic scale to facilitate comparison across orders of magnitude. The database contains 1,136 TFs, 751,590 cohesin-associated regulatory sites, and 15,195 genes. Based on these entities, Cohesin-GRN comprises 61,222,502 TF–site associations and 2,228,634 site–gene associations, which together give rise to 12,312,034 inferred TF– gene regulatory relationships. By integrating TF–site and site–gene links, the database further enumerates 275,616,774 TF–site–gene regulatory paths, representing distinct regulatory routes through which TFs can influence gene expression via different regulatory sites. Notably, the number of TF–site– gene paths is substantially higher than that of direct TF–gene associations, highlighting the pervasive use of multiple regulatory sites in TF-mediated gene regulation.

**Figure 2.**
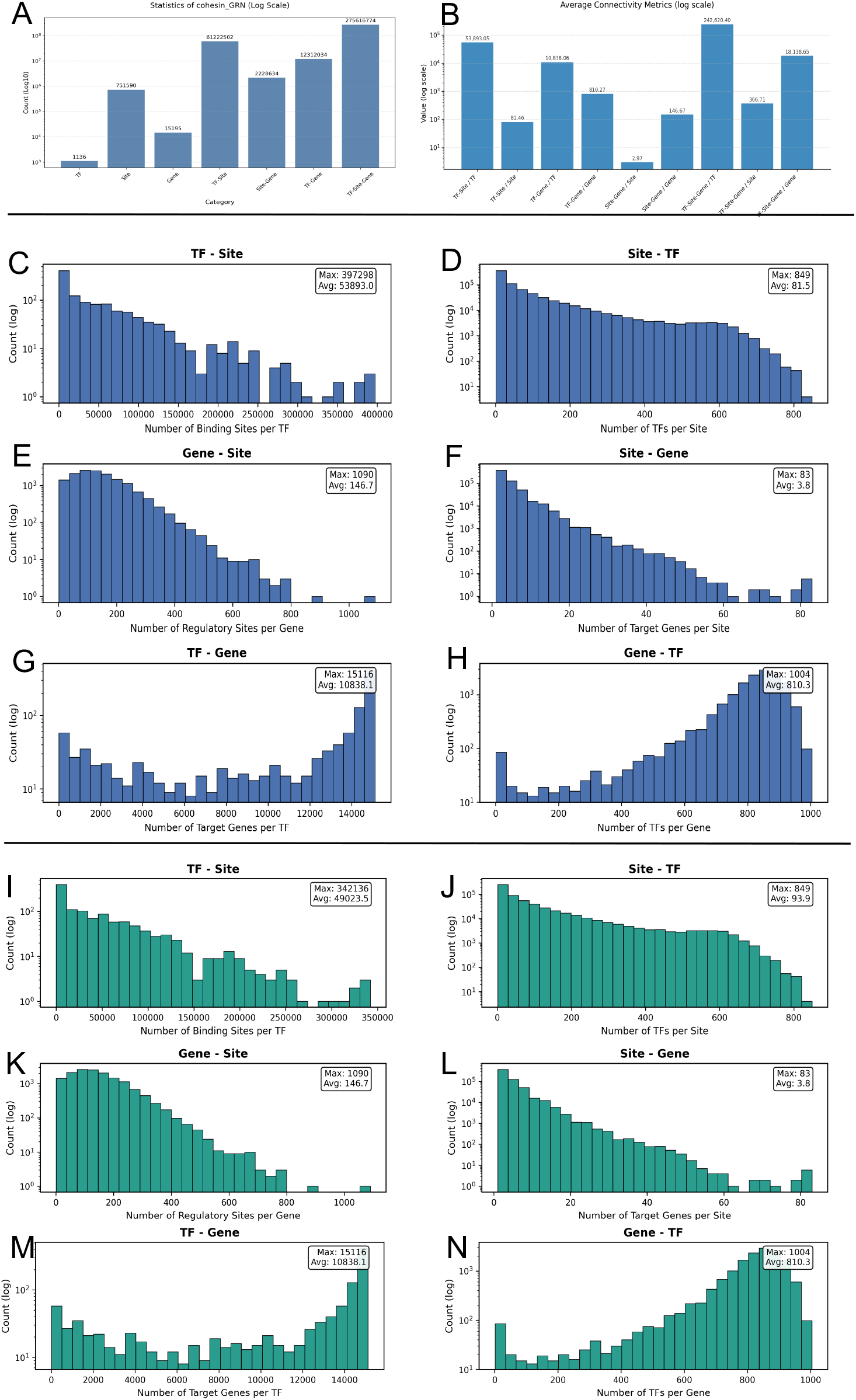
Statistical overview of regulatory information included in Cohesin-GRN. Summary of the scale, connectivity, and distribution of TF–site, site–gene, TF–gene, and TF–site–gene regulatory relationships in the cohesin-centered gene regulatory network.

**Figure 2B** illustrates the average connectivity metrics of TFs, regulatory sites, and genes across different layers of GRN. On average, each TF is associated with 53,893 regulatory sites, whereas each site is bound by approximately 81 TFs, indicating a highly many-to-many TF–site interaction structure. At the TF–gene level, each TF regulates an average of 10,838 genes, while each gene is regulated by approximately 810 TFs, reflecting extensive combinatorial regulation. In contrast, site–gene connectivity is markedly more constrained. Each regulatory site is linked to an average of only 2.97 target genes, representing the lowest connectivity among all relationship types and highlighting the relatively high specificity of regulatory sites for gene regulation. Conversely, each gene is associated with approximately 146 regulatory sites, indicating that genes typically integrate signals from multiple distinct regulatory elements. Integration of TF–site and site–gene relationships further expands the regulatory landscape at the TF–site–gene level. On average, each TF participates in approximately 242,620 TF–site–gene regulatory paths, each site participates in about 367 such paths, and each gene is involved in approximately 18,139 paths. Together, these connectivity patterns underscore the complexity of site-aware gene regulation, in which TFs exert combinatorial control over gene expression through large numbers of distinct regulatory sites.

### Distribution of regulatory relationships

The distributions of regulatory relationships across different layers of the cohesin-centered GRN are shown in **Figure 2C–H**. At the TF–site level (**Figure 2C**), most TFs are associated with a moderate number of regulatory sites, with an average of 53,893 sites per TF, whereas a small subset of TFs bind exceptionally large numbers of sites, reaching up to 397,298 sites. At the site–TF level (**Figure 2D**), most regulatory sites are bound by relatively few TFs (average 81.5), although a limited number of sites are targeted by hundreds of TFs, with a maximum of 849. At the gene–site level (**Figure 2E**), most genes are associated with a limited number of regulatory sites (average 146.7), while a small fraction of genes are linked to substantially larger regulatory neighborhoods, with up to 1,089 sites. For the site–gene distribution (**Figure 2F**), most regulatory sites regulate only a few genes, with an average of 3.8 target genes per site and a maximum of 83. TF–gene relationships further illustrate extensive combinatorial regulation. As shown in **Figure 2G**, individual TFs regulate highly variable numbers of target genes, with an average of 10,838 genes per TF and a maximum of 15,116. The distribution is bimodal, with relatively low frequencies in the intermediate range, indicating that most TFs either regulate a large set of genes (reflecting broad or conserved regulatory roles) or target a more restricted subset of genes with higher specificity. Despite binding tens of thousands of regulatory sites, each TF regulates a much smaller set of target genes (53,893 sites versus 10,838 genes on average), reflecting convergence of multiple regulatory sites onto the same genes. **Figure 2H** depicts the distribution of TFs regulating each gene. The distribution is strongly skewed toward high values, indicating that virtually no gene is regulated by a single TF. Instead, most genes are co-regulated by large numbers of TFs, with the majority falling between approximately 800 and 1,000 regulators. On average, each gene is regulated by 810.3 TFs, and the maximum number of TFs regulating a single gene reaches 1,004.

We next examined the distributions of regulatory relationships after restricting the network to TF– site–gene tripartite regulatory paths (**Figure 2I–N**). At the TF–site level (**Figure 2I**), TFs involved in tripartite regulation are associated with fewer regulatory sites compared with the full TF–site network shown in **Figure 2C**, with an average of 49,023 sites per TF. This reduction reflects the filtering effect imposed by requiring sites to participate in complete TF–site–gene paths. In contrast, sites retained in the tripartite network tend to be bound by more TFs. As shown in **Figure 2J**, each site involved in TF– site–gene relationships is regulated by an average of 93 TFs, which is higher than the corresponding value observed in **Figure 2D**. This enrichment suggests that sites participating in tripartite regulation are more likely to serve as regulatory hubs. The distributions of site–gene relationships (**Figure 2K and 2L**) closely mirror those observed in the full network (**Figure 2E and 2F**). Similarly, TF–gene connectivity patterns in the tripartite network (**Figure 2M and 2N**) are highly consistent with those in the full TF– gene network (**Figure 2G and 2H**). This consistency is expected, as TF–gene regulatory relationships are inferred through the integration of TF–site and site–gene associations within the tripartite framework.

## Usage of search and browse Cohesin-GRN

Figure 3. illustrates the usage workflow and result presentation of the cohesin-GRN web platform. **Figure 2A-B** show the single-search and multi-search interfaces, respectively, which support flexible querying of TFs, genes, and genomic regions either individually or in combination. As an example, the single-search mode is used to query the transcription factor ZNF391, the gene FNDC10, and the genomic interval chr1:1,000,000–1,500,000. The query results are summarized in **Figure 3C**, which provides a statistical overview of the returned regulatory relationships in the specified region. The summary includes the numbers of TFs, genes, and unique regulatory sites identified after deduplication, as well as the numbers of TF–site, site–gene, TF–gene, and TF–site–gene relationships. In this example, TF, Gene, and Site represent the numbers of TFs, genes, and unique regulatory sites identified within the queried genomic region, respectively. TF–Site indicates the number of cohesin sites identified with ZNF391 motif located in the specified interval, whereas Site–Gene denotes the number of regulatory sites through which FNDC10 is regulated in the same region. TF–Gene represents the number of inferred TF–gene regulatory pairs; because a single TF and a single gene are queried and only one regulatory relationship exists between them within the specified interval, this value equals one. Importantly, TF–Site–Gene captures distinct regulatory paths in which a TF regulates a target gene through different regulatory sites. In this case, a value of one indicates that ZNF391 regulates FNDC10 through a single regulatory site within the chr1:1,000,000–1,500,000 region.

**Figure 3.**
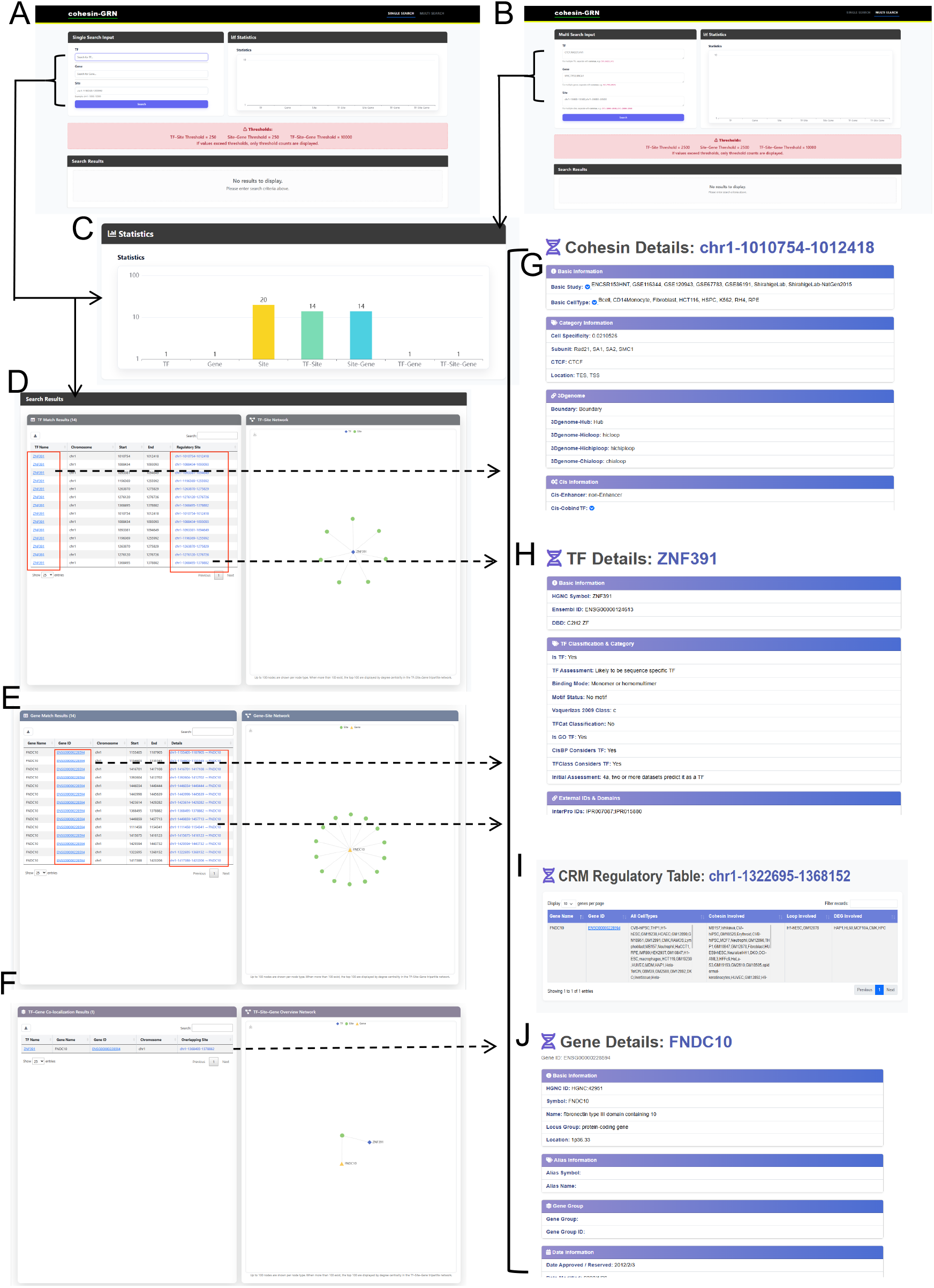
Usage examples and interface screenshots of the Cohesin-GRN web platform. Demonstration of single-search and multi-search workflows, result summaries, interactive tables, and network visualizations for exploring TF–site–gene regulatory relationships.

**Figure 3D–J** present the search results returned by the Cohesin-GRN web platform. To ensure system performance and maintain visual clarity, display thresholds are applied to the query outputs: for TF–site and site–gene relationships, only the top 250 records are shown, whereas up to 10,000 records are displayed for TF–site–gene relationships. The results are organized into three sections corresponding to TF–site, site–gene, and TF–site–gene regulatory relationships. Each section includes both a tabular view and a corresponding force-directed network visualization. In the network views, node selection is based on degree centrality calculated from the TF–site–gene regulatory network, and the top 100 nodes ranked by degree centrality are displayed to highlight potentially important regulatory hubs. The force-directed network graphs support interactive exploration: hovering over a node highlights its neighboring nodes and displays detailed annotation information, while hovering over an edge reveals the specific regulatory relationship between the connected entities.

**Figure 3D** illustrates the TF–site search results in Cohesin-GRN. The left panel presents a results table listing TFs and their associated regulatory sites, with TF names provided as clickable links to access detailed annotation pages. An example TF detail page is shown in **Figure 3H**, which summarizes basic information, functional categories, and relevant annotations. Regulatory sites are likewise interactive, and selecting a site opens a dedicated annotation page (**Figure 3G**) displaying information such as TFs binding to the selected site. The right panel of **Figure 3D** shows the corresponding force-directed network graph, providing an intuitive overview of TF–site regulatory relationships. **Figure 3E** displays the site–gene search results. The results table lists genes, gene identifiers, associated regulatory sites, and the corresponding site–gene regulatory pairs. Clicking on a gene name opens the gene detail page (**Figure 3J**), while selecting a specific site–gene pair enables inspection of detailed site–gene information in tabular form (**Figure 3I**). The accompanying force-directed network graph visualizes regulatory relationships between sites and genes. **Figure 3F** presents the integrated TF–site–gene results, which constitute the core output of the Cohesin-GRN platform. The results table summarizes TFs, target genes, gene identifiers, and the regulatory sites linking each TF–gene pair, while the corresponding network visualization highlights how TFs regulate target genes through specific regulatory sites.

## Discussions

In this study, we constructed a cohesin-centric GRN database that explicitly incorporates regulatory site information into network representation. Compared with conventional TF–gene GRN models, which typically ignore regulatory sites, the inclusion of cohesin-associated CREs enables a more detailed and mechanistically informative view of transcriptional regulation. From the perspective of GRN analysis [19], site-aware network representations provide deeper insights into regulatory architecture than direct TF–gene relationships alone. From the perspective of cohesin biology [10], organizing cohesin-associated regulation in a network-based framework substantially improves the efficiency of exploring complex regulatory mechanisms mediated by cohesin

Although cohesin is treated as a unified class of regulatory sites in the current version of Cohesin-GRN, cohesin binding sites are known to play distinct functional roles in genome regulation [16]. In particular, cohesin co-occupied by CTCF are more likely to participate in chromatin structural organization, whereas non-CTCF cohesin sites are more frequently involved in CRE-mediated transcriptional regulation [15]. In the present implementation, all cohesin binding sites are included without explicit functional stratification. Future versions of Cohesin-GRN will incorporate functional classification of cohesin sites to distinguish structural and regulatory roles more precisely. Another limitation is that regulatory relationships in the TF–site–gene network are defined as unweighted edges, with all interactions treated as contributing equally. This simplification may complicate biological interpretation, particularly given the large number of regulatory sites associated with each gene (on average 146 sites per gene). Incorporating quantitative weights, such as genomic distance between sites and genes [30] or H3K27ac intensity, would enable more refined modeling of regulatory influence and improve interpretability of GRN analyses. Moreover, TF–site relationships in the current Cohesin-GRN are inferred primarily from motif-based predictions, and TF–gene regulation is obtained by merging TF– site and site–gene relationships. Future incorporation of more advanced regulatory inference methods is expected to improve both the precision and interpretability of the resulting GRN.

CohesinDB has served as a resource for studying cohesin biology in human cells. As CohesinDB continues to expand with additional datasets and methodological updates, the introduction of Cohesin-GRN module provides a novel framework for investigating cohesin-mediated transcriptional regulation and gene regulatory networks.

## Author contributions

Jiankang Wang conceived and designed the study. Jinyu Ding and Jiankang Wang constructed the database, developed the web interface, and wrote the manuscript. Haoping Chen, Yanqiang Fu, Wenze Gao, and Zhixiong Fang contributed to data analysis. All authors reviewed and approved the final manuscript.

## Acknowledgement

This work was supported by the National Natural Science Foundation of China (32400526, W2523104), the Changsha Municipal Natural Science Foundation (kq2402059), Hunan Provincial Natural Science Foundation of China (2024JJ6131), the Open Research Fund of the State Key Laboratory of Gene Expression (SKLGE-KF-2025002), the Guangdong Basic and Applied Basic Research Foundation (2023A1515110873, 2025A1515012822), and the Fundamental Research Funds for the Central Universities (541109030087), all awarded to Jiankang Wang.

